# SNP discovery and analysis of genetic diversity between commercial crossbred and indigenous chickens from three different agro-ecological zones using DArT-Seq technology

**DOI:** 10.1101/2023.02.23.529829

**Authors:** Kwaku Adomako, Selorm Sovi, Bismark Kyei, Jacob Alhassan Hamidu, Oscar Simon Olympio, Samuel E. Aggrey

## Abstract

Indigenous and commercial chickens have developed unique adaptations to their environments, which may include nutrition, pathogens, and thermal stress. Besides, environmental pressures and artificial selection have generated significant genome-wide divergence in chickens, as those selection pressures contribute a considerable evolutionary force to phenotypic and genotypic differentiation. Herein, we determined genomic diversity of indigenous chickens from semi-deciduous rainforest (SDR), coastal savannah (CC) and Guinea savannah (GS) agro-ecological zones (AEZs) in Ghana and commercial crossbreds reared at the Kwame Nkrumah University of Science and Technology (KNUST). We generated SNP markers from 82 chickens (62 indigenous chicken ecotypes and 26 commercial crossbred ecotype) using DArT-Seq technology. A total of 85,396 SNP markers were generated and after filtering the data, 58,353 markers were used to study genetic diversity and population structure analyses. Polymorphism information content (PIC) values ranged from 0.0 to 0.5, with 21,285 SNP markers (35%) being in the lowest PIC value range (0 to 0.15) while 13,511 (21%) were in the highest PIC value range (0.45 to 0.50). Between the KNUST population and the indigenous ecotypes, pairwise F_ST_ values were estimated to be 0.105 between CS, 0.096 between SDF, and 0.133 between GS. Furthermore, PCA analysis showed that the CC, SDF and GS chickens clustered together and are genetically distant from the commercial crossbred. We herein show that chickens from the AEZs studied can be considered as one population. However, due the abundance of agro-byproducts in the SDR compared to the CS and GS, chickens from the SDR AEZ had better growth compared to their counterparts. It is suggested that the genetic diversity within the local ecotypes could form the basis for genetic improvement.

## Introduction

Chicken is one of the most common domesticated species, as it plays a key role in agricultural and biomedical research fields. Backyard, small-scale and commercial chicken production contribute significantly to human livelihood and food security of poor households [1, 2]. In addition, their impact goes beyond the provision of food, cash income, and employment, as they also serve as a means of capital accumulation and are valued in religious and sociocultural lives of most people [3, 4]. Chickens are found on all habitable continents, agro-ecological zones (AEZs), and their morphology and traits vary, which may affect their genetic makeup [5, 6]. According to some studies, a challenging environment can shape the genomic landscape that underlies a population’s adaptation to climate and resources [7, 8]. Li et al. [9] stated that environmental pressures and artificial selection can generate significant genome-wide divergence in chickens, as those selection pressures contribute a considerable evolutionary force to phenotypic and genotypic differentiation. Understanding the genetic diversity and differentiation of chickens commonly found in Ghana would provide the basis for genetic conservation and/or genetic improvement.

In the last two decades, Diversity Arrays Technology (DArT) has produced a high-throughput marker technique called DArTseq based on genotyping-by-sequencing (GBS) to sequence the most informative representations of genomic DNA samples [10, 11]. DArTseq™ produces a fairly simple and accurate source of genetic data, which can also be used further for additional population analysis [12]. Furthermore, it integrates the genome complexity reduction method [13] with next-generation sequencing (NGS) approach. The technology merges high-throughput processing of DNA to produce complexity-reduced genome representations using combinations of restriction enzymes to select the optimal fraction of the genome for sequencing with rigorous analytical procedures [12]. This application has been successfully established in several plant [14, 15] and animal species [16, 17]. In addition, DArTseq has been applied to a wide range of species and applications, including the study of inter- and intra-specific genetic diversity and relationships, genetic mapping, genome-wide association studies, and genomic selection [18–21]. DArTseq produces single nucleotide polymorphism (SNP) markers that have been successfully applied for genetic structure analysis in several animal species [17, 22, 23].

The present study aimed to characterize the phenotypic diversity between chickens from three different AEZs and commercial crossbred chickens; and identify the genomic diversity of chickens within three AEZs and commercial crossbreds using SNP markers. A comprehensive and deep understanding of the genome underpinnings of the indigenous and commercial crossbreds could reveal the genetic diversity and population structure of these chickens in different AEZs. They can serve as the basis for genetic conservation and/or genetic improvement.

## Materials and methods

### Ethics statement

All the procedures for this experiment were conducted under a protocol approved by the Institutional Ethical Committee at Kwame Nkrumah University of Science and Technology (KNUST), Ghana.

### Sampling of the indigenous and commercial crossbred

We randomly selected 104 birds for this experiment. The sampled birds were comprised of the indigenous chicken ecotypes and commercial crossbreds. A total of 72 indigenous chickens were randomly selected, comprising 36 males and 36 females. We sampled 12 males and 12 females from each AEZ (Guinea Savannah, Semi-Deciduous Rainforest, and the Coastal Savannah agro-ecological). In addition, 16 male and 16 female commercial crossbreds were sampled from a crossbred population housed at KNUST [24]. The crossbreds constitute an exotic Lohmann breed mated randomly with indigenous ecotypes. Since then, the various first filial males have been backcrossed to the Lohmann female for 7 successive generations. Hereafter, the Lohman x indigenous ecotype would be referred to as KNUST.

### Description of the ecological zones

The Guinea Savannah zone lies between longitude 100 1’ W and latitudes 100 3’ and 110 10’ N. The climate of the location is normally dry with a unimodal rainfall pattern that starts from April to October. The average annual rainfall ranges from 800 to 1200 millimeters. The dry season runs from November to March/April, with the hottest temperatures occurring near the conclusion of the season. The average temperature is 32°C and the average relative humidity (RH) is 33%.

The Semi-Deciduous Rainforest is found in the country’s middle belt and is located between longitude 2.250 W and 0.150 W and latitude7.460 N and 5.500 N. The average yearly rainfall is between 1200 and 1600 millimeters. The zone experiences two rainy seasons: the major season runs from March to July, and the minor season runs from September to November. RH varies from 97 and 20% through to the morning and late afternoon, respectively, with an average temperature of about 27°C.

The climate of the Coastal Savannah is a tropical type. Temperatures are regularly high, with very little difference during the year. The yearly average temperature is 26.80°C, with monthly temperatures ranging from 24°C in August to 29°C in March. RH is generally high in the area and varies from 65% in the midday to 95% at night. The bimodal rainfall pattern in the zone gives rise to two rainy growing seasons with an average yearly rainfall of 800mm. The major rainy season runs from May to mid-July, and the minor rainy season runs from mid-August to mid-October.

### Phenotypic traits

Phenotypic traits measured on all birds at 36 weeks of age were: body weight (BW), body length (BL), body width (BWd) and shank length (SL).

### Collection of blood sample and genomic DNA extraction

Whole blood was sampled from all chickens. Total genomic DNA was extracted from chicken whole blood using the QIAGEN® Kit (QIAGEN. Valencia, CA, USA) at the Molecular Genetics Laboratory of the Department of Animal Science, University of Ghana, Legon. The DNA concentration in the collected samples was determined using a Life Technologies Qubit 3.0 fluorometer. The Qubit was used for evaluating DNA quality prior to next-generation sequencing (dsDNA). According to the manufacturer’s procedures, a Qubit dsDNA BR (wide range, 2 to 1000 ng) Assay Kit and Qubit dsDNA HS (high sensitivity, 0.2 to 100 Nanograms) Assay Kit were used with a Qubit 3.0 fluorometer; a sample volume of 1 μL was added to 199 μL of a Qubit working solution.

### Genotyping chicken accessions using SNP markers

DNA samples from 90 chickens were sent to the Biosciences Eastern and Central Africa (BecA)-ILRI Hub, Nairobi, Kenya (https://www.ilri.org/research/programs/beca-ilri-hub) for genotyping by sequencing (GBS) using DArTseq™ technology on Illumina HiSeq2500. Sequencing data was obtained from 62 indigenous chicken ecotypes and 26 commercial crossbred individuals. The sequences were aligned to the Chicken Reference Sequence, Ggal 5 to identify chromosomes and positions. SilicoDArT SNP identification was performed with MSQ 0.6.6. Only readings that aligned to a single unique site of the genome were considered for SNP finding. SNP markers were scored ‘1’ for presence, and ‘0’ for absence, and ‘-’ for failure to score. To calculate the marker data’s reproducibility, two technical replicates of the DNA samples were genotyped.

## Statistical Analysis

### Analysis of phenotypic data

Two-way ANOVA was conducted on the phenotypic data using the 12th edition Genstat statistical software[25], significance was established at p<0.05, and differences between treatment means were differentiated using the Tukey’s Studentized Range Test.

### Analysis of sequencing data

#### Quality analysis of SNP marker data

The “R” software (R x64 v4.0.2) was used to test the markers for reproducibility (%), call rate (%), polymorphism information content (PIC), and one ratio. The proportion of technical replicate assay pairs was used to score reproducibility. The call rate was calculated using the percentage of samples with a score of ‘0’ or ‘1’ to determine the outcome of reading the marker sequence across the samples. Polymorphic information content is the degree of diversity of the marker in the population and shows the usefulness of the marker for linkage analysis. One ratio constitutes the proportion of the samples for which genotype scores equaled ‘1’. The PIC refers to the marker’s degree of diversity in the population, and it demonstrated the marker’s usefulness in linkage analysis. The fraction of samples with genotype scores of ‘1’ is represented by one ratio.

#### Population structure and genetic diversity

Analysis of population structure was conducted in R software version 3.3.1. Nei’s distances were computed to estimate genetic differences in proportion to time. Nei’s genetic distances were used to calculate Analysis of Molecular Variance (AMOVA). The AMOVA, as implemented by the “poppr.amova” function, was used to determine variance within and between the chicken populations. The “stampp F_ST_” function was used to calculate genetic differentiation (F_ST_) across all populations’ pairs. Using principal component analysis (PCA) with the “find. Clusters” function in R, sampling sites investigated the genetic structure, population differentiation, and AEZ depending on the available allelic frequency. Clustering was done using the “aboot” function from the “poppr” package. Discriminant Analysis of Principal Components (DAPC) analysis was employed to identify and describe the number of clusters of genetically related individuals. DAPC plot emphasizes between-group variation as well as within-group variation. The contributions of alleles to the DAPC groupings aid in identifying regions of the genome that drive genetic divergence between populations [26]. A dendrogram was generated for the population differences in neck type, and AEZs and individuals were categorized into a predetermined number of population groups/clusters (K).

The software program STRUCTURE was used to apply a Bayesian methodology based on the genotypes of the individuals collected [27]. Using a probabilistic approach, individuals in the dataset were assigned to K (unknown) populations. Individuals are allocated membership to all of the distinct clusters (number of clusters = K), where the sum of the probabilities belong to a population equals one (1), and K varies between runs of the program. To ensure a random starting point for the algorithm, STRUCTURE was run with 106 iterations and a burn-in period of 10,000 iterations. To ensure that the findings were consistent, the runs were performed 20 times for 2 greater than (>) K less than (<) 10. A FastStructure, an alternative to Structure, was created specifically for huge SNP datasets [28], like the admixture, ancestry model, which assumes neither polyploid nor dominant data and allows individual birds to have mixed ancestry. This is modeled by assuming that a certain individual has acquired some portion of its genomic DNA from ancestors in population K. This allowed for a graphical representation of the data to be created in order to distinguish between different groups.

## Results

### Phenotypic characterization

The phenotypic characteristics of the chickens studied are presented in Table 1. The chickens from the Semi-deciduous rainforest were heavier and broader (P<0.05) when compared with their counterparts from the Coastal Guinea Savannah. Body length and shank length were not significant (P>0.05) among the chickens from the three AEZs. There was sexual dimorphism among the commercial crossbreds located at KNUST. The males were had higher body weight, body width, body length and shank length (P<0.05) when compared with the females.

**Table 1.** Performance of indigenous chickens from the Coastal Savannah (CS), Semi-deciduous rainforest (CDF), Guinea Savannah (GS) and commercial crossbred chickens located at Kwame Nkrumah University of Science and Technology (KNUST) in Ghana at 36 weeks of age

| Location | Body weight (g) | Body length (cm) | Body width (cm) | Shank length (cm) |
| --- | --- | --- | --- | --- |
| CS | 846<sup>b</sup> | 35.08 | 7.63<sup>ab</sup> | 8.00 |
| SDR | 1,425<sup>a</sup> | 34.38 | 7.85<sup>a</sup> | 8.21 |
| GS | 962<sup>b</sup> | 35.21 | 7.21<sup>b</sup> | 7.88 |
| SEM | 113 | 0.94 | 0.23 | 0.27 |
| <sup>a-b</sup>Pr>F | 0.001 | 0.645 | 0.026 | 0.495 |
| KNUST |  |  |  |  |
| Male | 2,437<sup>a</sup> | 53.70<sup>a</sup> | 14.81<sup>a</sup> | 11.81<sup>a</sup> |
| Female | 1,862<sup>b</sup> | 39.02<sup>b</sup> | 11.14<sup>b</sup> | 9.05<sup>b</sup> |
| SEM | 0.022 | 0.099 | 0.278 | 0.092 |
| Pr>F | <0.001 | 0<.001 | <0.001 | <0.001 |

<sup>a-b</sup>Means in the same column with different superscript are significantly different at $P<0.05$

### Discovery and quality analysis of SNP markers

A total of 85,396 SNP markers were generated. After removing missing data scaffolds with no specific chromosomal assignment, minor allele frequency of less than 5%, sex chromosomes, and chromosomes with no positions, 58,353 SNP markers remained for further analysis. The quality of SNP markers was assessed using call rate and PIC. Approximately 70% of SNP markers were more than 95% reproducible and remained within 90% to 95% range. The call rate of 58,353 SNP markers varied from 30% to 100%, with 70% showing ≥ 80% call rate and the remaining markers showing < 80% call rate (Fig 1A). The PIC values for the SNP markers ranged from 0.00 to 0.50. Around 21,285 SNP markers (35%) were in the lowest PIC value range (0 to 0.15), while 13,511 (21%) were in the highest PIC value range (0.45 to 0.50) (Fig 1B). The majority of SNP markers, 48,644 (83%) were found on chromosome 1 to 15, with the remaining markers, 9,709 (17%) found on chromosome 16 to 33 (Fig 1C). Moreover, few SNP markers were found on chromosome 16, 31, and 32, with no SNP markers on chromosomes 29 and 30. These findings indicate that SNP markers are of high quality and may have a significant impact on the chickens’ genome.

**Fig 1.**
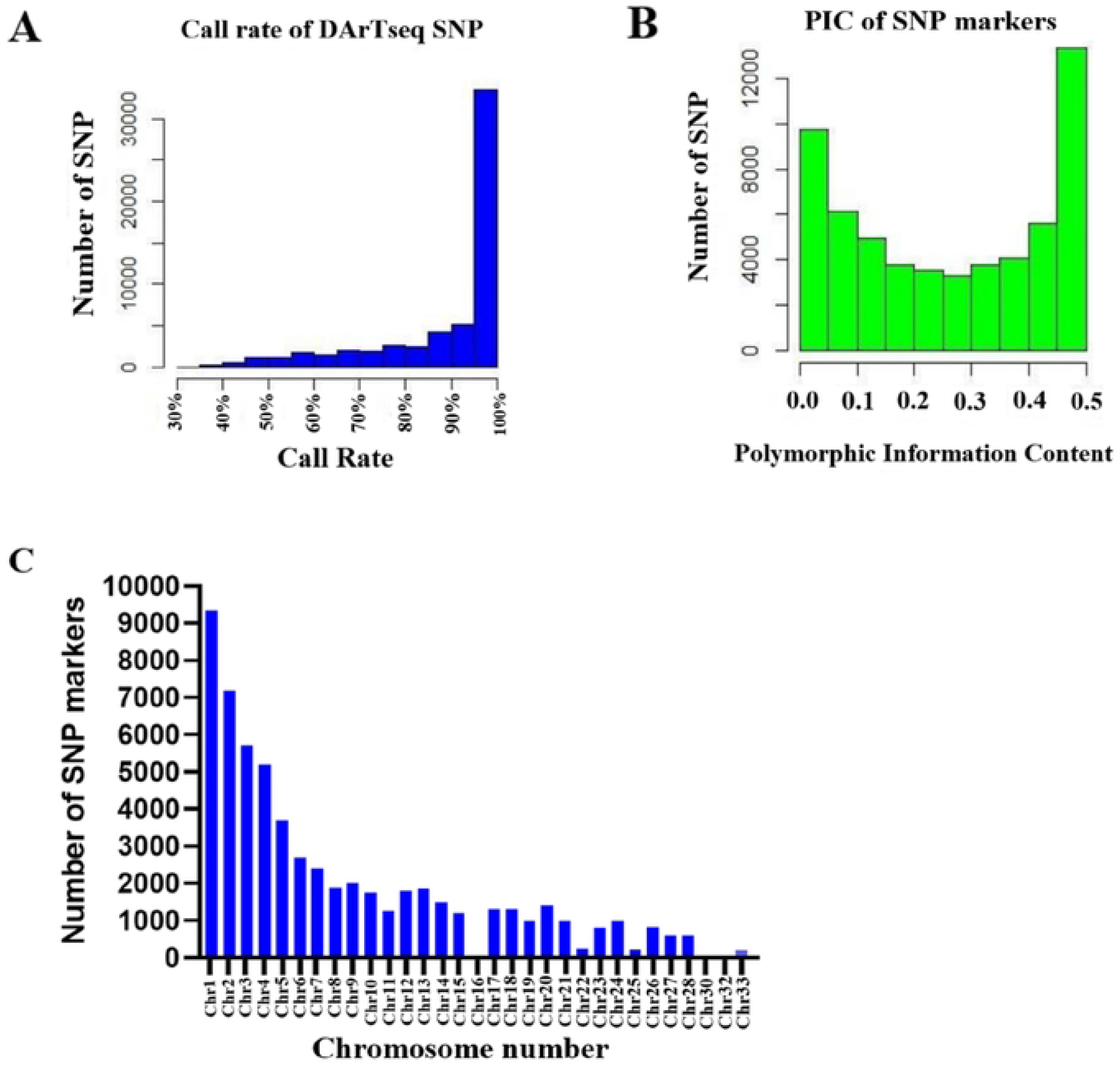
Qualities and characterization of SNP markers in chicken genotype. (A) Call rate of SNP markers tested in chicken genotypes. (B) Frequency distribution of PIC values for SNP markers. (C) Distribution of SNP markers on the chromosome of chicken.

### Genetic diversity among population structure

F_ST_ of the AEZs/ecotypes based on SNPs was performed to obtain the total genetic variance contained in a subpopulation relative to the total genetic variance (Table 2). Between KNUST lines and the indigenous ecotypes, pairwise F_ST_ values were estimated as 0.105 between Coastal Savannah lines, 0.096 between Semi-Deciduous Rainforest lines, and 0.133 between Guinea Savannah lines. Besides, the highest FST value obtained from pairwise comparison of the three local ecotypes was 0.031 between Guinea Savannah and Semi-Deciduous Rainforest lines; and the lowest F_ST_ value obtained was 0.015 between Semi-Deciduous Rainforest lines and Coastal Savannah lines. We found that the highest level of genetic differentiation was between commercial crossbred chicken breeds from KNUST and Guinea Savannah zone populations (F_ST_ = 0.133). These findings indicate that there is high genetic structure between commercial crossbreds from KNUST and indigenous lines from the three AEZs.

**Table 2.**
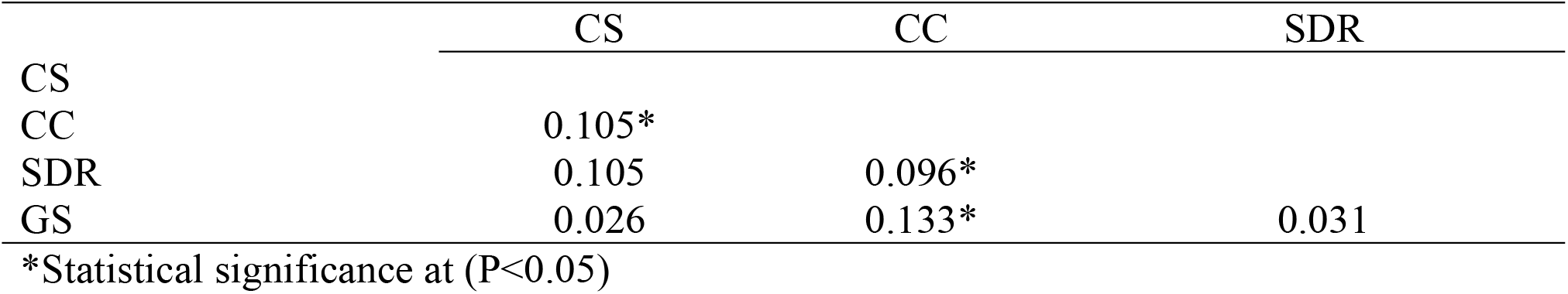
Pairwise F_ST_ among chickens from Coastal Savannah (CS), Commercial crossbreds (CC), Semi-deciduous Rainforest (SDF) and Guinea Savannah (GS) based on SNP data

An agglomerative hierarchical cluster analysis was performed on SNP markers using Nei’s genetic distances to study the genetic relationships between the chicken genotypes. The Nei’s genetic distance measures the extent of genetic variation that exists between species. Herein, chickens from the various AEZs, including the commercial crossbreds from KNUST, were considered. The Nei’s genetic distance values were lower than 0.068 in all populations (Table 3). on the SNP calls. The commercial crossbreds and the Coastal Savannah chickens had a proportionate differentiation of 0.057, and that of the commercial crossbreds and Semi-Deciduous Rainforest was 0.053, and that between the commercial crossbreds and Guinea Savannah was 0.068. In a similar manner, the genetic distance between the Coastal Savannah and Semi-Deciduous Rainforest lines was 0.019, Guinea Savannah and Semi-Deciduous Rainforest was 0.025, as well as 0.021 between the Coastal Savannah and Guinea Savannah. These findings showed that there is more genetic overlap between the commercial crossbreds and Guinea Savannah chickens, followed by the commercial crossbreds and the Coastal Savannah chickens.

**Table 3.** Nei’s genetic distance among chickens from Coastal Savannah (CS), Commercial crossbreds (CC), Semi-deciduous Rainforest (SDF) and Guinea Savannah (GS) based on SNP data

|  | CS | CC | SDR |
| --- | --- | --- | --- |
| CS |  |  |  |
| CC | 0.057* |  |  |
| SDR | 0.019 | 0.053* |  |
| GS | 0.021 | 0.068* | 0.025 |

\*Statistical significance at ( $P < 0.05$ )

### Dendrogram analysis

The dendrogram generated was based on the Nei’s genetic distance [29] among individuals within the chicken (Fig 2). Besides, the clustering based on the SNP markers produced two main clusters of related populations or similar origin. Cluster 1 consisted of all the commercial crossbreds. The clustering within the commercial crossbreds had various degrees of similarity, ranging from a high similarity (100%) to a lower similarity index (68.9%). Two individual birds from the commercial crossbreds identified as “KNU 13” and “KNU 5” are the least diverged individuals among this cluster, with Nei’s genetic distance of 0.09 and a high similarity index of 100%.

**Fig 2.**
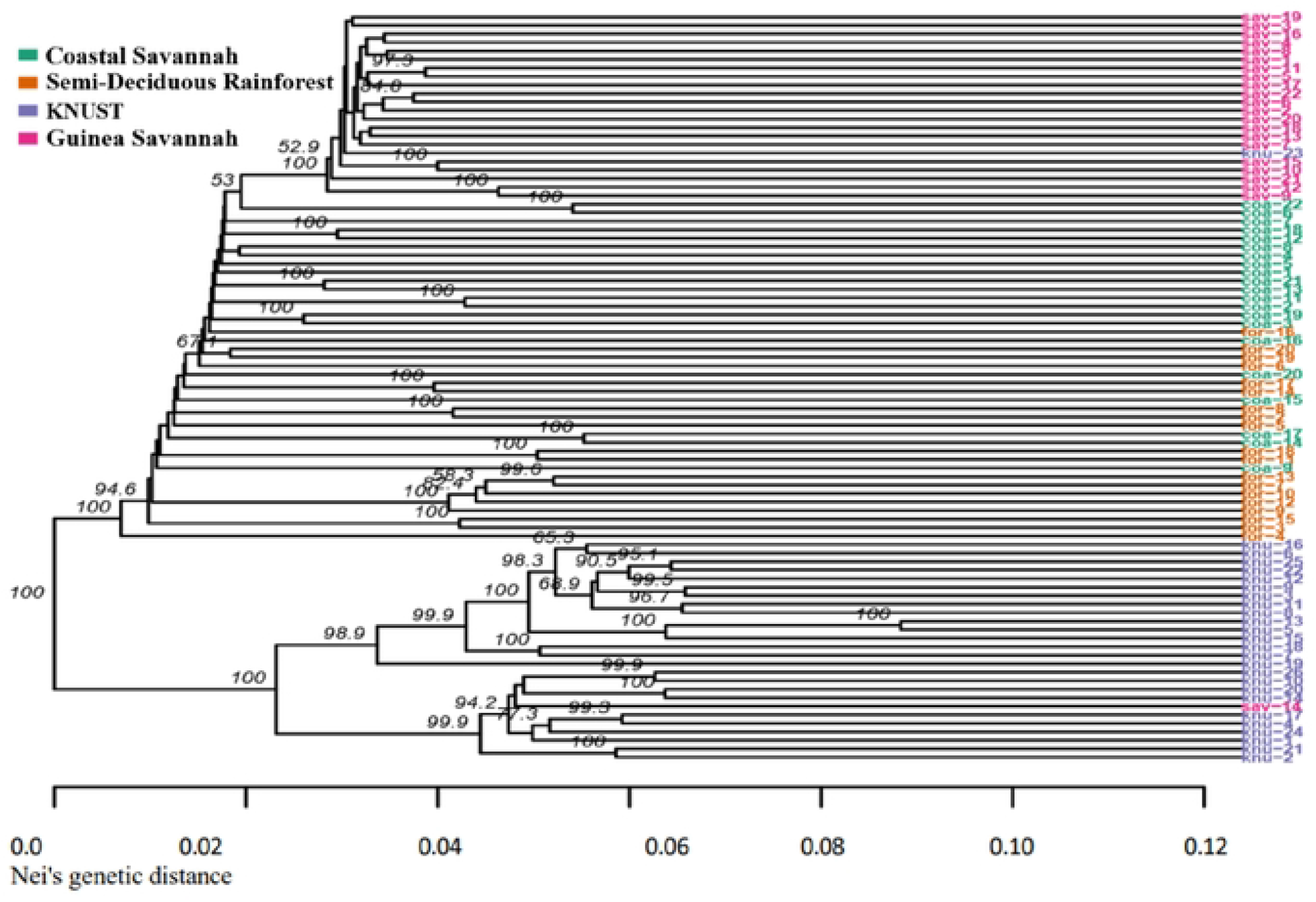
Dendrogram showing relatedness among chicken genotypes based on SNP markers Detection of number of clusters and principal component analysis.

The indigenous ecotypes were considered as one cluster however, the Guinea Savannah population formed a sub-group within the cluster. From the current data, the SNP markers data showed a shorter Nei’s genetic distance ranging from 0.00 to 0.12. While the Semi-Deciduous Rainforest and Coastal Savannah populations tended to cluster together, it appeared that Semi-Deciduous Rainforest, Coastal Savannah, and Guinea Savannah populations shared most SNP alleles and were less diverged. However, they also shared a degree of similarity ranging from high similarity (100%) to low similarity index (52.9%). However, the commercial crossbreds based at KNUST had a wide range of diversity as compared to the other populations.

To confirm the separation based on commercial crossbred chickens (KNUST) and chickens from the three ecological zones, the data was subjected to PCA plot to show the presence of potential clusters of populations, with principal components jointly explaining the total genetic variance in the chicken populations using the SNP data. Score scatter plots showed the differentiation and similarities among the chickens (Fig 3). The first principal component highlighted the genetic variation between the chicken populations. We separated the cluster comprising chicken populations belonging to the Semi-Deciduous Rainforest, Guinea Savannah, and Coastal Savannah zones from the commercial crossbred chickens, which had extended to two main different clusters. The commercial crossbred population from KNUST was more dispersed and appeared not to share much similarity with the chicken ecotypes for the three AEZs. This analysis shows that chickens from KNUST have a diverse genetic composition when compared to indigenous chickens from the ecotypes.

**Fig 3.**
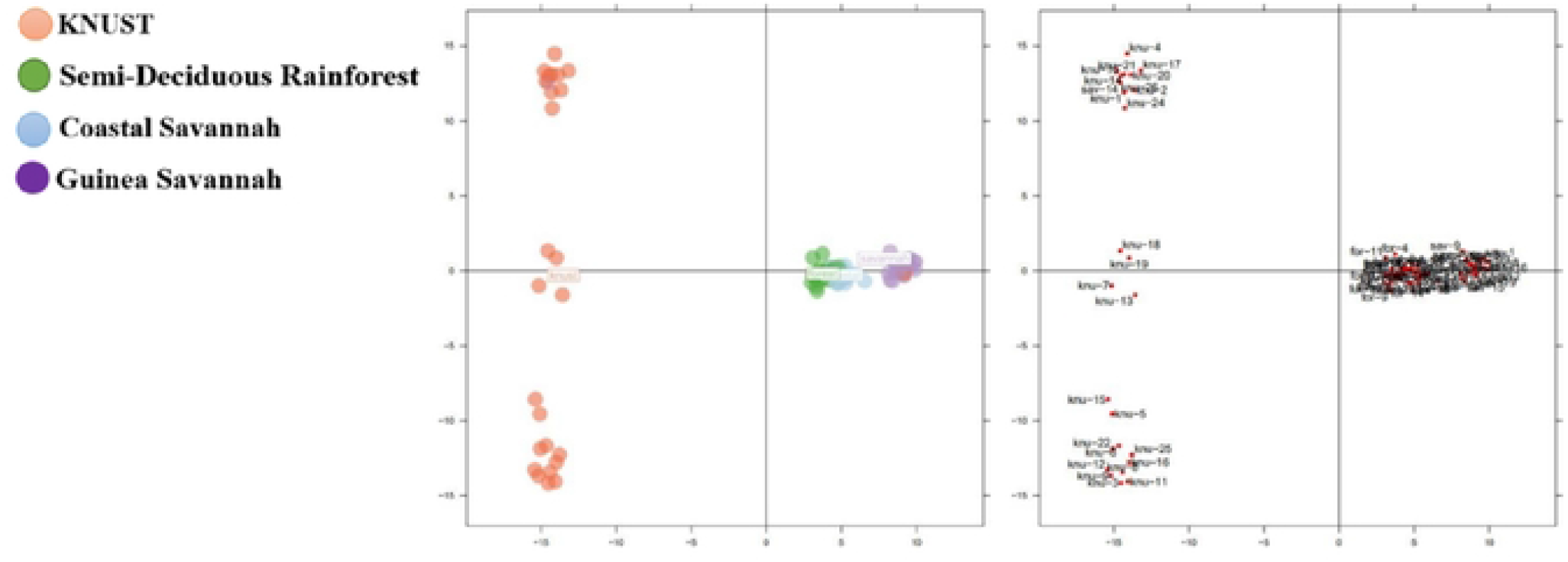
Principal component analysis of the chicken genotype based on SNP markers.

### Analysis of molecular variance

Analysis of molecular variance (AMOVA) was conducted to partition the molecular diversity among and within populations for SNP marker data. As presented in Table 4, the AMOVA revealed that the highest level of variation (82%) was within the samples’ variance. However, types within the population had the lowest variation (1.062%). Also, the variation between the chicken populations was 6.9%. Results from the population structure analysis are shown in Fig 4. The analysis indicates that the best grouping of the population was two (K=2). A total of 26 individuals, mainly from the commercial crossbreds (KNUST) were entirely grouped in population 1 (deep blue), while 22 chickens mainly from Guinea Savannah were grouped in population 2 (light blue). Forty chickens mostly from Coastal Savannah and Semi-Deciduous Rainforest ecotypes were in the admixture group. However, the larger proportion of the individuals from the admixture group were from the Guinea Savannah ecotype.

**Fig 4.**
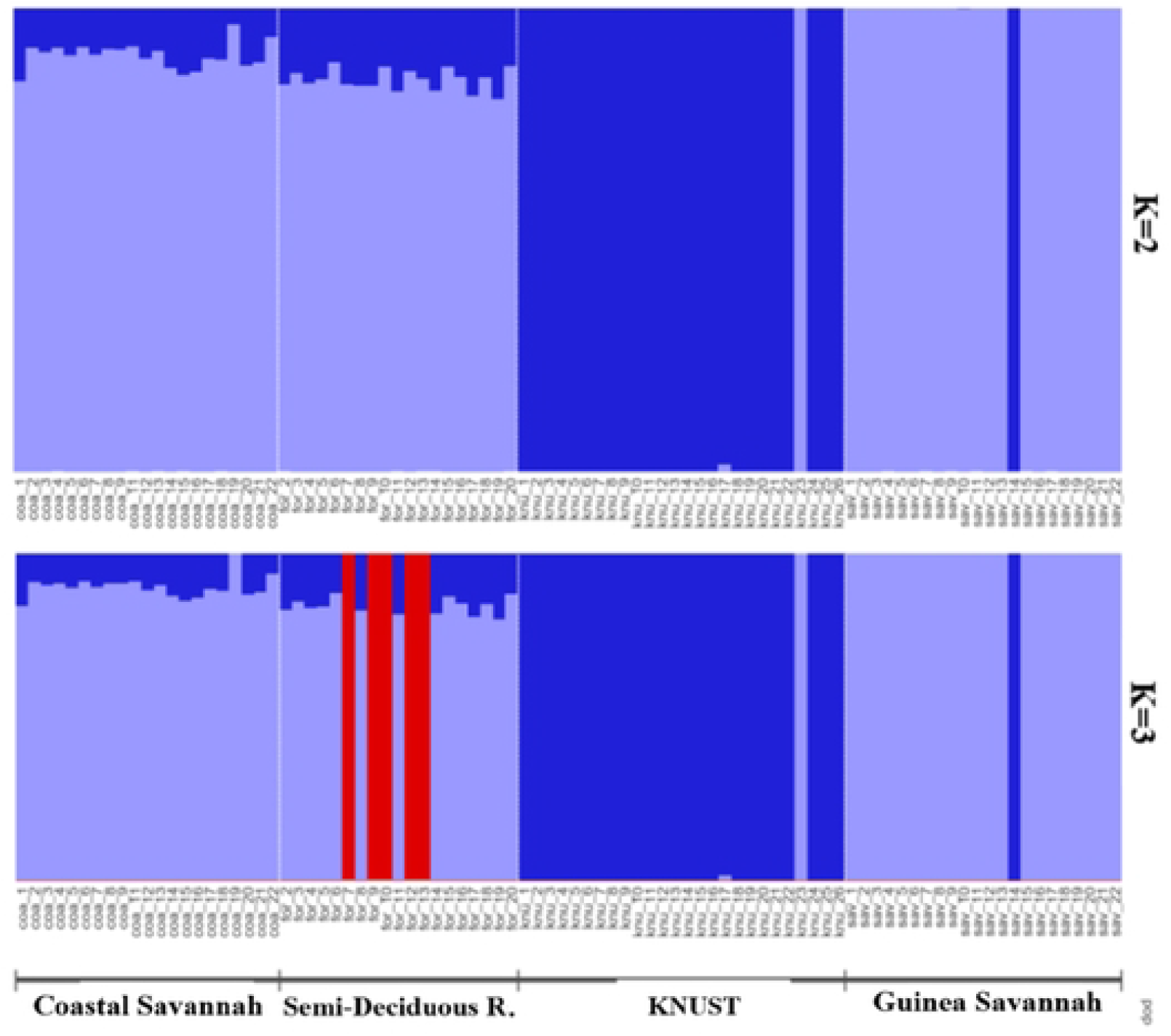
A structure bar plot showing the assignment of indigenous chicken and commercial crossbred (KNUST) populations. Each vertical bar represents one population and different color within each bar indicate admixture. (K = 2).

**Table 4.** Analysis of molecular variance based on SNP marker data.

| Table 4. Analysis of molecular variance based on SNP marker data |  |  |  |  |  |
| --- | --- | --- | --- | --- | --- |
| Source of variation | DF | SS | MS | Sigma | Variance (%) |
| Between chicken population | 4 | 22,777 | 5,694 | 112 | 6.92 |
| Between type with chicken population | 9 | 16,723 | 1,858 | 17 | 1.06 |
| Between samples with type | 74 | 122,409 | 1,564 | 159 | 9.76 |
| Within samples | 88 | 117,655 | 1,337 | 1,337 | 82.26 |
| Total | 175 | 279,564 | 10,453 | 1,625 | 100.00 |

## Discussion

The current study aimed at investigating some of the phenotypic differences among chickens located in three AEZs (Coastal Savannah, Semi-deciduous Rainforest and Guinea Savannah) and commercial crossbreds located at KNUST. Among the local ecotypes, chickens from the Semi-deciduous rainforest was the heaviest compared to their counterparts in the coastal Savannah and the Guinea Savannah. Most local chicken ecotypes are reared in the backyard and rely primarily on scavenging for their nutrition. Household scraps and agro-byproducts are their main source of nutrition. The yearly average rainfall of Semi-deciduous Rainforest (1,200-1,600 mm) is adequate for the cultivation of large-scale plantation crops such as cocoa, oil palm and lemon, as well as, annual crops (maize, cassava and plantain) [30]. By products from such crops, insects and worms for a bulk of their diet. The abundant agro-byproducts in the Semi-deciduous rainforest contribute to the better growth of chicken ecotypes in that AEZ compared to their counterparts in the Savannahs. The Semi-deciduous rainforest may be the most suitable environment to raise indigenous chickens whose sustenance depend primarily on scavenging.

Current studies indicate that the use of SNP markers has significant role to play in determining genetic diversity and population structure in animal species, such as in amphibians [31], grasshoppers [32], fishes [33] and lobsters [34] and chickens [22, 35]. For example, SNP dataset is known to provide more reliable inferences of patterns of genetic structure and diversity than a typical microsatellite dataset in Liberian amphibians [31]. In addition, Dementieva et al. used SNP markers to analyzed genetic diversity of different chicken breeds [35]. Therefore, studying the genetic diversity and population structure of chickens from three AEZs and KNUST using SNP markers, may contribute to the understanding the diversity and structure of chickens reared in different AEZs in Ghana. In the current study, we found that there is high genetic diversity and population structure between the commercial crossbreds and indigenous chickens across the AEZs.

The AEZs are directly and indirectly affect the survival [36], productivity [37, 38], and adaptation [39] of animals. An animal of high genetic potential may perform poorly in a resource poor AEZ [40]. Currently, data on phenotypes, genetic diversity and population structure among chickens in different AEZs in Ghana is scant. Through the application of DArTseq genotyping, a total of 58,353 reads of SNP markers were discovered and used for genetic diversity characterization in indigenous and commercial chickens. SNPs form the basis for large DArTseq genotyping platforms [41] and can also contribute to the forthcoming version of the chicken genome. The SNP markers used in this study showed an average genotype call rate of 90%, showing that the quality of the SNPs was high. The PIC value reveals information about the diversity. PIC values can be categorized into three: high (0.40 to 0.5), moderate (0.10 to 0.25), and low (0 to 0.10) [42, 43]. In the current study, 35% of SNP markers were in the lowest PIC value range (0 to 0.15), while 21% were in the highest PIC value range (0.45 to 0.50), with an average value of 0.26. Thus, about 67% of the SNP markers discovered showed moderate to high polymorphism and could be used for other genome wide studies. The DArTseq markers (SNPs) used in this study had an average PIC value of 0.26, which was lower than what was observed in Korean and Iranian native chicken using microsatellite markers [44, 45]. Moreover, when compared to some studies in plant species where DArTseq markers were used, the average PIC value was similar, and in some cases higher than what was observed in these plants. For example, in durum wheat, it was 0.265, while in a study of watermelon it was 0.13 [46, 47]. This indicates that the SNP markers from the study were significantly informative for genetic diversity and genome wide association studies in chickens.

The pairwise FST statistic among population is used to measure the genetic distance among populations [48]. The low FST values among the indigenous ecotypes indicate low genetic differentiation among these populations, which means they might have a common origin. However, there were high F_ST_ values among the crossbreds at KNUST and the indigenous chicken ecotypes. Considering the Nei genetic index, we observed a higher genetic diversity between the crossbreds located at KNUST and the indigenous groups, which could be attributed to the differences between the exotic parental types of the crossbreds and the indigenous parental type. The crossbreds located at KNUST and the Coastal Savannah chickens had a proportionate differentiation of 0.057 suggesting that these two chicken populations may have some overlap. their DNA segments. This suggest that the crossbred may share ancestry with some of the chickens in the Coastal Savannah. The genetic differentiation between the Coastal and Guinea Savannah and that of Coastal and Semi-Deciduous Rainforest had the lowest estimated distance. This implies that there is less genetic diversity between the populations with shorter genetic distances, which in turn mirrors the population history with a likely common origin.

A dendrogram demonstrates attribute distances and relationships of similarities among a group of entities. In the current study, we constructed a dendrogram between commercial crossbred chickens (KNUST) and indigenous chickens from different AEZs based on significant SNP markers. The SNP dendrogram mainly consisted of two clusters. The dendrogram based on genetic distances in Fig 2 shows that there is a clear difference between KNUST (cluster 1) and indigenous chickens from the AEZs (cluster 2). This clearly corroborate the FST results that shows that the commercial crossbreds cluster differently from the indigenous ecotypes.

Additionally, we used the PCA method to ascertain differences and similarities among the different groups of chickens [49]. We ascertained that chickens from the three AEZs were located in the same areas of the loading plot and clustered together, indicating good repeatability of the test and a putative common origin. However, the commercial crossbreds located at KNUST were clearly separated from the indigenous chickens in the loading plot. We used AMOVA to examine the genetic variation among and within the different chicken groups [50]. The current study revealed that there was 6.92% genetic variation between the chicken population and that most of the variation came from variation within samples among populations. Similar outcome was observed in four Thai indigenous chickens [51]. The current study showed arrangement in structure output at K=2 which suggests a high genetic differentiation among the studied chicken populations and once again, the chickens from the Guinea Savannah, Semi-Deciduous Rainforest, and Coastal Savannah AEZs clustered together and are farther away genetically from the commercial crossbred chickens at KNUST.

## Conclusion

In the current study, we investigated the genetic structure and variability in chickens from three AEZs and commercial crossbreds raised at the Kwame Nkrumah University of Science and Technology (KNUST) using SNP markers. Using population genetics parameters, FST, Nei’s genetic index, dendrogram and PCA analysis, we demonstrated that the chickens from the Semi-Deciduous Rainforest, Coastal Savannah and Guinea Savannah may have a common genetic origin and can be considered as one genetic population. These local ecotypes are different in structure and are genetically distant from the commercial crossbred chickens reared in KNUST. However, due to abundant agro-byproduct resources in the Semi-Deciduous Rainforest, the chickens in this AEZ had better growth than the chickens reared in the Savannahs. The AMOVA analysis also showed that there is ample diversity within sample genetic variability suggesting that, these indigenous populations may not be under immediate threat of inbreeding and can play a significant role in future genetic improvement programs.

## Acknowledgments

The authors are grateful to the Kwame Nkrumah University of Science and Technology Research Fund (KReF) for funding the research. We are also grateful to Dr. Alexander W. Kena for his assistance in the data analysis. Last but not the least, we would like to thank Prof. B. B. Kayang for allowing us to use his molecular laboratory for the genomic DNA extraction.

## Authors contributions

Conceptualization: Kwaku Adomako, Samuel E. Aggrey

Funding acquisition: Kwaku Adomako.

Investigation: Kwaku Adomako, Selorm Sovi

Methodology: Kwaku Adomako, Selorm Sovi.

Data Collection: Kwaku Adomako, Selorm Sovi, Bismark Kyei.

Formal analysis: Jacob Alhassan Hamidu, Oscar Simon Olympio, Bismark Kyei.

Supervision: Kwaku Adomako, Oscar Simon Olympio.

Writing-original draft: Kwaku Adomako, Selorm Sovi, Bismark Kyei.

Writing-review and editing: Kwaku Adomako, Samuel E. Aggrey

